# IL-17-producing CD8^+^ T cells differ in phenotype, metabolic profile and cytokine production to their CD4^+^ Th17 counterparts

**DOI:** 10.64898/2026.01.25.701586

**Authors:** Jessica Anania, Yaqing Sera Ou, Zachary Lim, Angus Moffat, Sarah Dimeloe, Emily Gwyer Findlay

**Author notes:** corresponding author –. these authors contributed equally (Yaqing Sera Ou, Zachary Lim).

## Abstract

Th17 cells (CD4^+^ T cells producing IL-17) are important for clearance of fungal infections and play a critical role in the development and exacerbation of numerous autoimmune diseases. Their differentiation, signalling pathways, cytokine production and metabolism are now well-characterised. As well as CD4^+^ T cells, CD8^+^ cells also produce IL-17 family members, and these have been named Tc17 cells. However, much less is known about their development, signalling or metabolism compared to their CD4+ counterparts. Here, we performed a series of *in vitro* and *in vivo* analyses of Tc17 cells as well as computational analysis of the published *Tabula muris* dataset, comparing Tc17 to IL-17^−^ CD8^+^ T cells and to Th17 cells. We show that murine Tc17 cells are generated in the presence of TGF-β and IL-6, and that cells produced by these culture conditions substantially reflect Tc17 cells seen *in vivo;* that is, with high expression of PD1, CD6, ICOS and CD161. Tc17 cells show phenotypic and functional differences to their Th17 counterparts, with increased production of IL-2 and IL-22 as well as an increased tendency to produce IL-17F as well as IL-17A. They show a more glycolytic profile than Th17 cells, with lowered mitochondrial membrane potential. This divergent phenotype and cytokine production suggests differential roles *in vivo* for these two cells.

## Introduction

Ever since their discovery 20 years ago ^1–3^, Th17 cells (CD4+ T cells producing IL-17) have been extensively characterised. Their importance and unique roles in immunity are now clear - they are essential for defence against fungal infections and play critical roles in host defence and homeostasis at mucosal sites in particular. The differentiation, signalling and metabolism of Th17 cells is now well phenotyped (reviewed in ^4,5^). Their induction is driven by IL-6 and TGF-β signalling, resulting in phosphorylation of STAT3, activation of SMAD family members, the RORγt transcription factor and the aryl hydrocarbon receptor, and production of IL-17 cytokines as well as others including IL-21. As well as their key roles in host defence, Th17 cells drive and exacerbate autoimmune disease such as rheumatoid arthritis, psoriasis and Multiple Sclerosis (reviewed in ^6^).

CD8^+^ T cells have also been observed to produce IL-17, and these cells have been named Tc17 (cytotoxic CD8^+^ T cells producing IL-17). They are most commonly found in mucosal tissue ^7–10^. They are up-regulated during inflammatory disease, and their importance in autoimmune conditions has been noted. While Tc17 cells play a key role in viral, fungal and bacterial immunoprotection ^11^, they have also been implicated in various inflammatory diseases including psoriasis, Multiple Sclerosis, neuromyelitis optica, and inflammatory bowel disease ^9,12–17^, as well as many types of cancer ^18–21^. They are suggested to be directly pro-carcinogenic.

Despite this importance, extensive characterisation of Tc17 induction and differentiation has not been performed and many questions remain about the differentiation and regulation of these cells. It is unclear whether their signalling or metabolism differ to their Th17 counterparts, or whether they play different roles in autoimmune disease induction.

Here, we set out to characterise murine Tc17 cells *in vivo* and *in vitro*. We established optimum differentiation conditions, defined their cell surface markers and transcription factor expression, characterised their cytokine production, and determined their metabolic differences compared to both Th17 cells as well as to Tc1 cells.

## Results

### Generation of Tc17 cells requires TGF-β

Firstly, we set out to establish a culture model which generated consistent numbers of murine IL-17^+^ CD8^+^ T cells. There is significant variation in the methodology used by previous papers in this topic. It was originally shown that TGFβ and IL-6 stimulate CD8^+^ cells to differentiate into noncytotoxic, IL-17-producing cells ^22^. Since then, however, various combinations of IL-1β, IL-6, TGF-β and IL-23 have been used *in vitro* ^23^, with many papers ^7,24–27^ stating that IL-1β is required for consistent generation.

To determine the optimum method for Tc17 generation, we compared published methods over 48 and 96 hours of culture. Consistent production of IL-17A by CD8^+^ T cells was observed following incubation with IL-23, IL-6 and TGF-β (Fig1A,B), the same culture conditions which drive the highest production from CD4+ T cells^28^. In contrast to other papers, we did not observe substantial expression of IL-17A in the presence of IL-23 and IL-1β (Fig.1A,B). In all conditions, inclusion of αCD28 antibodies enhanced production.

**FIGURE 1:**
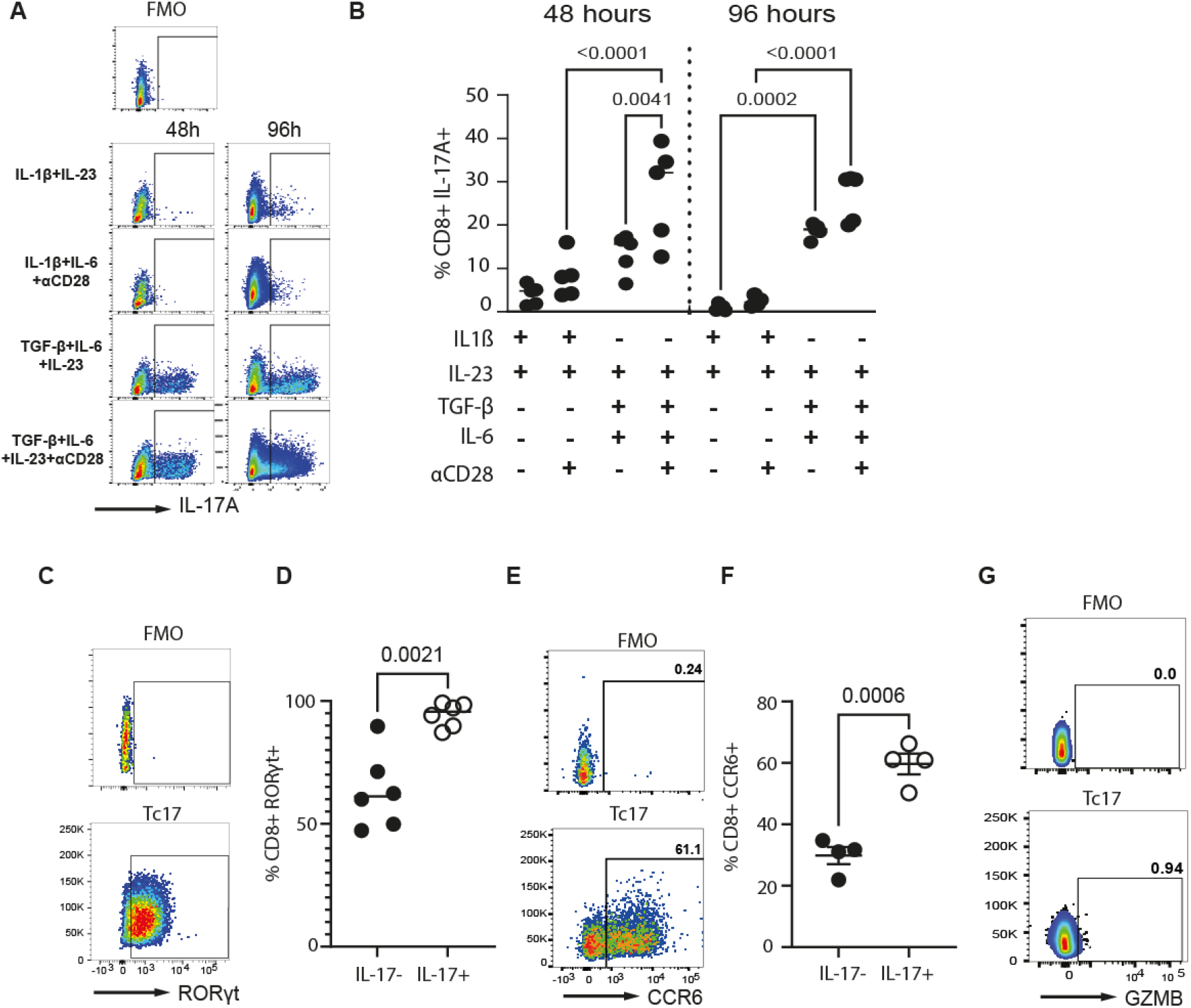
Generation of Tc17 cells requires TGFβ. Splenocytes isolated from C57BL/6J mice were cultured for 48 or 96 hours in the presence of a combination of Th 17-d riving cytokines. (A, B) Expression of IL-17A by CD8+T cells in culture was detected by flow cytometry. All IL-17A+ cells were (C,D) RORγt+ and the majority were (E,F) CCR6+. All were (G) granzyme B negative. Data shown are individual mice used in separate experiments with line at median. Statistical significance was determined using (B) a two way A NOVA with Tu key’s multiple comparisons test and (D,F) paired Student’s t test. N values: B-5; D - 6; F - 4.

To confirm that these cells were truly Tc17 cells, we established that they expressed the master transcription factor for Tc17 cells, RORγt (Fig.1C,D). They also had significantly enhanced expression of CCR6, which is strongly associated with IL-17+ cells ^10,29,30^) (Fig.1E,F), and were negative for granzyme B, the characteristic mediator produced by classical cytotoxic CD8^+^ cells which is absent from Tc17 cells ^31^ (Fig.1G).

### Analysis of gene expression in Tc17 cells *in vivo*

Having generated large numbers of Tc17 cells in culture, we started by understanding which signalling pathways are triggered in CD8^+^ T cells that drive their differentiation into Tc17 cells vs cytotoxic Tc1 cells, and whether their gene expression is substantially similar to Th17 cells. To answer this question, we first carried out analysis of the *Tabula muris* dataset ^32^. This dataset contains single cell RNA sequencing analysis of 356,213 cells from 20 mouse organs. We selected for analysis CD8^+^ IL-17^−^, CD8^+^ Il-17^+^ and CD4^+^ Il-17^+^ cells; the tissue origin of selected cells was predominantly spleen, thymus, lung and bone marrow. Following normalisation and standardisation, gene expression profiles of three cell populations were compared using the Scanpy package.

Tc17 cells were, as expected, substantially divergent in gene expression to their IL-17-counterparts, with an increase noted in the receptors for IL-17-driving cytokines IL-23 and TGF-β, as well as in IL17-associated transcription factors (*Rora, Rorc, Id2*). Compared to IL-17^−^ CD8^+^ T cells, there were also observed to be large increases in expression of co-stimulatory molecules and other markers which reflect a high state of activation, such as *Cd5, Cd6, Cd69*, and *Icos*.

Some striking differences were noted in IL-17^+^ cells in this dataset compared to IL-17^−^ cells, which were consistent between the Th17 and Tc17 subsets. These included an increase in *Batf* and large increases in *Runx3, Maf*, and *Ikzf3*. This strong expression of *Maf* ties in with observed increase in the gene coding for PD1, *Pdcd1*, as Maf has previously been shown to be induced by TGF-β and lead to PD1 expression in CD8+ T cells ^33^.

However, there were also clear transcriptional differences between Tc17 and Th17 cells. Of note, *Il22* and *Id2* were strongly increased in Tc17 cells compared to Th17; in contrast, *Ccr4* and *Ccr6* were much lower in Tc17 cells compared to Th17.

### Cytokine production by Tc17 cells

Next, we examined cytokine patterns in Tc17 compared to Th17 cells. In addition to IL-17A, production of IL-17F was also observed, with the same pattern of induction as IL-17A (that is, maximally produced following incubation with recombinant TGF-β, IL-6, IL-23 and anti-CD28 antibodies (Fig.3A,B). Interestingly, production of IL-17F was substantially higher than IL-17A in CD8^+^ T cells following culture, more so than is seen in Th17 cells (Fig.3C). This is in contrast to that observed for human cells^9^, in which IL-17A was the predominant family member produced. To understand this more, we looked at the ratio of production in *ex vivo* steady state splenic T cells. These produced more IL-17A than F, matching the human cells (Fig3D). Following induction in culture, as previously stated, Tc17 cells produced more IL-17F than IL-17A.

In addition, we assessed production of IFN-γ. In the absence of TGF-β, there was substantial production of IFN-γ, which was enhanced with CD28 stimulation (Fig.3E,F), indicating that cells produced with IL-1β and IL-23 maintain a cytotoxic Tc1 profile. However, the same culture conditions which generated the most IL-17 (that is, including TGF-β) also suppressed IFN-γ, confirming that these conditions are inducing Tc17 cells. At 96h, the IL-17A: IFN-γ ratio for these culture conditions was 6:1 (Fig.3G), while in all other conditions it was under 1, indicating that in the absence of TGF-β the CD8^+^ T cells continued to produce more IFN-γ than IL-17A.

We quantified GMCSF production; this is a hallmark cytokine of a particular subset of Th17 cells, designated pathogenic Th17 for their role in inducing mouse models of Multiple Sclerosis ^34^. Tc17 cells generated in culture expressed more GMCSF than IL17-CD8+ T cells, but this was extremely low, with on average 0.8% of cells producing it (Fig.3H). Following the identification of IL-21 and IL-22 as differentially regulated in Tc17 cells in the *Tabula muris* dataset, we also tested their production at the protein level. Flow cytometry data on cultured Tc17 cells reflected the patterns seen with scRNAseq data, with IL-21 not increased in Tc17 cells while IL-22 was increased. Finally, previous work has shown Tc17 cells produce TNF ^25^, and so we compared examined these cytokines too. Re-capitulating previous work, we found that Tc17 cells strongly express TNF, but so did IL-17^−^ cells. Interestingly, IL-2 was also noted to be very strongly up-regulated in Tc17 cells, in contrast to Th17 cells where it was absent.

### Tc17 cells show different surface marker expression compared to IL-17-CD8+ T cells and Th17 cells

We next analysed the surface marker pattern on Tc17 cells, to compare to both IL-17 negative cytotoxic CD8^+^ T cells and to CD4^+^ Th17 cells. It has previously been shown that human Tc17 cells can be identified by a pattern of high expression of CD6, CD69, CD39, PD1, and CD120b ^9^, but equivalent analysis has not been performed for murine cells. Murine CD8^+^ IL-17^−^ and IL-17A^+^ populations were therefore compared to evaluate whether Tc17 cells expressed a different pattern of cell surface markers to IL-17-cells, as a result of their differential gene expression patterns. Choice of markers was informed by the previously cited study and by the *Tabula muris* analysis in Figure 2.

**Figure 2:**
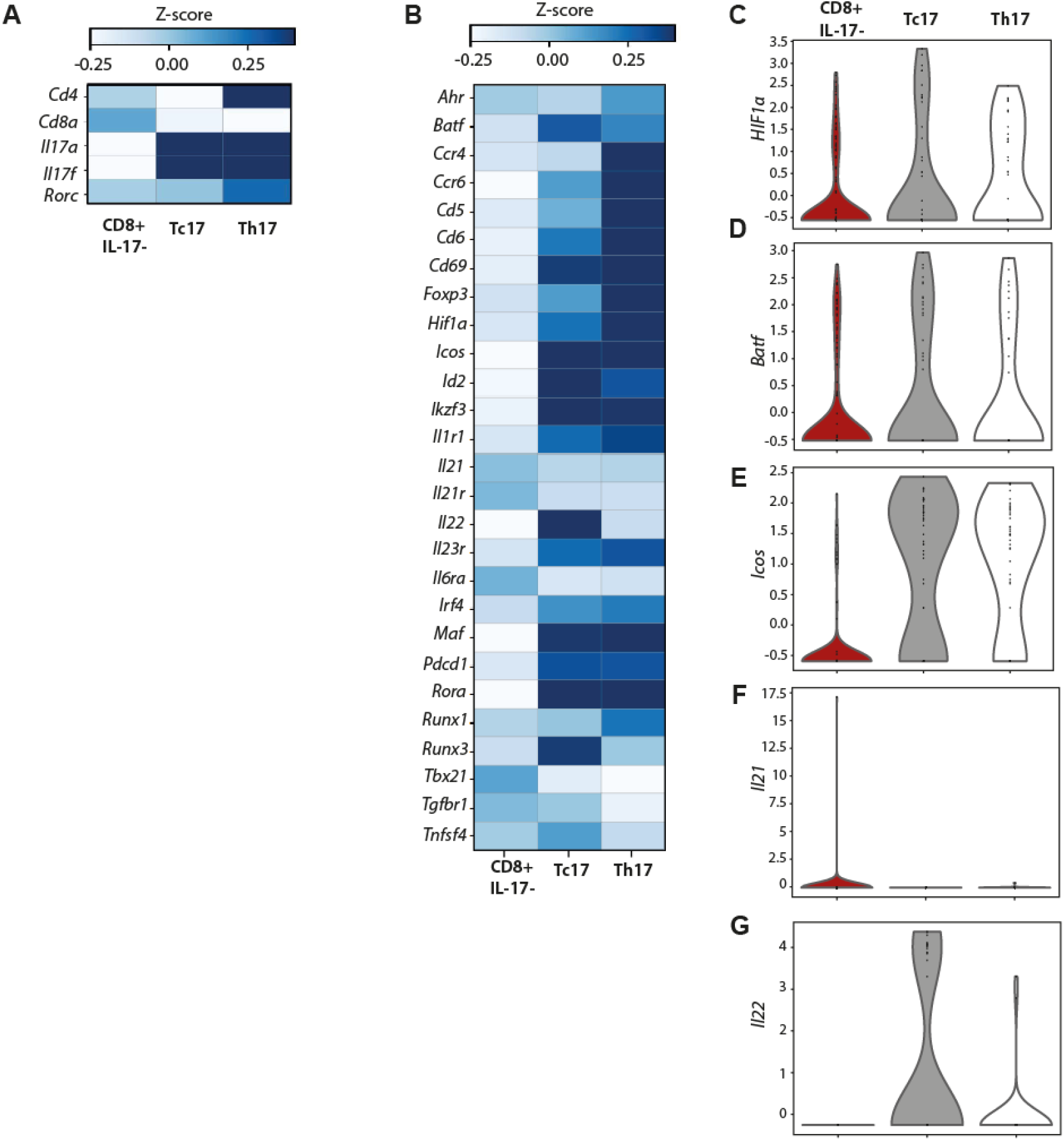
Tc17 cells *in vivo* have a distinct transcriptional profile compared to IL-17-CD8+ T cells or Th17 cells. Analysis of the Tabula muris dataset identified (A)Tc17 cells by their expression of CD8α and IL-17 cytokines. (B) the transcriptional profile of these cells wsa assessed and (C-G) high expression of *Hiflα, Batf, Icos* and *ll22*, and low expression of *ll21* noted.

**Figure 3:**
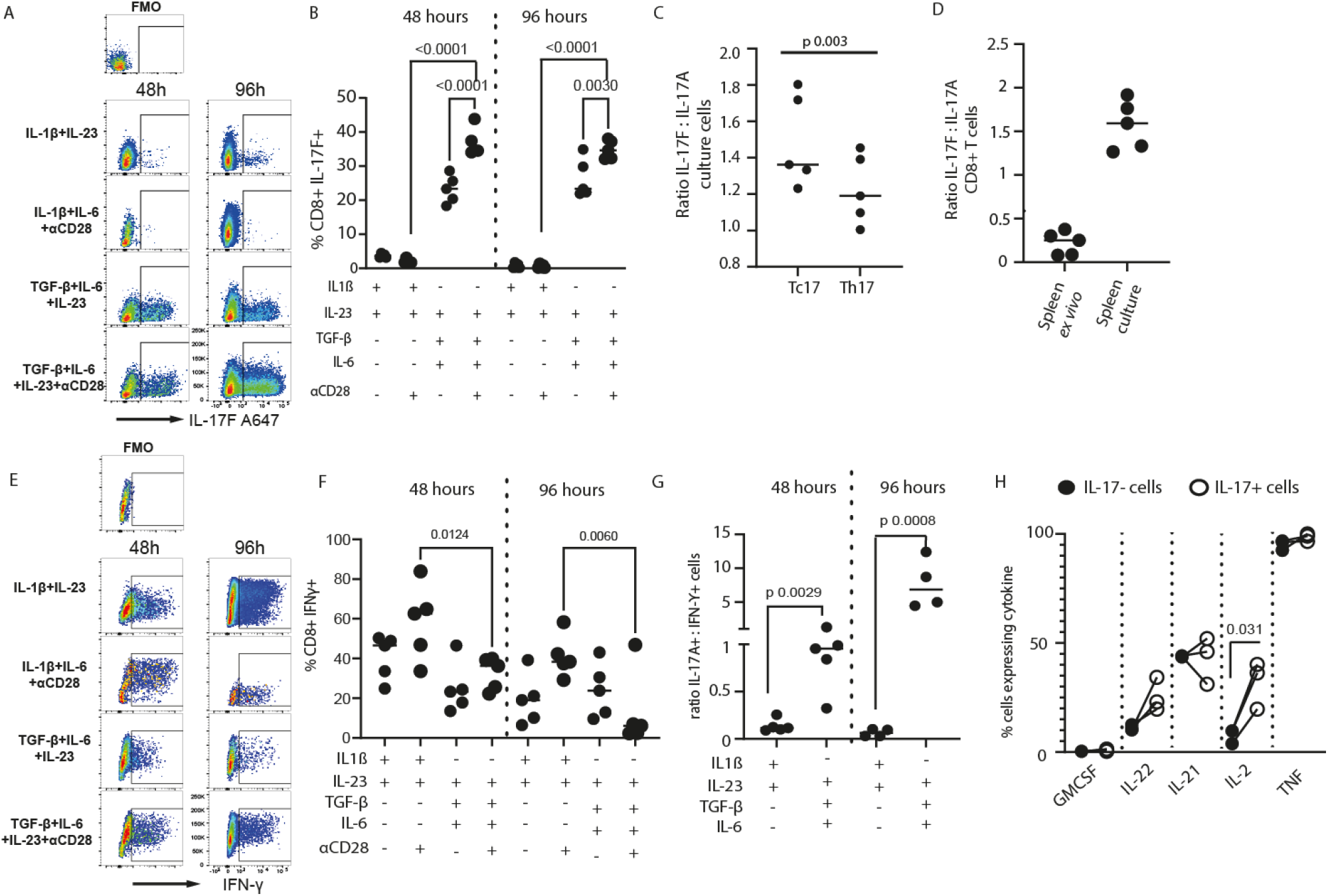
Cytokine production by murine Tel7 cells. Splenocytes isolated from C57BL/6J mice were cultured for 48 or 96 hours in the presence of Tc17-d riving cytokine combinations. Expression of (A,B) IL-17F was assessed by flow cytometry and (C) compared to IL-17A production. (C) IL-17F:A ratio was quantified in spleen CD8+T cells either from direct isolation ex vivo or following 48 hours in culture. (E-F) Expression of IFN-γ in cultured splenic CD8+T cells was quantified after 48 and 96 hours and (G) ratio of IL-17: IFN-γ was calculated. (H) Expression of other cytokine production by CD8+ IL 17+or IL-17-cells 48 hours of culture was quantified by flow cytometry. Data shown are individual mice with lines at median. Statistical significance was determined using (B, F) two-way ANOVA withTukey’s multiple comparisons test; (C) ratio paired Student’s t test; (G, H) paired Student’s t tests. N values = 3-5.

ICOS, CD6, CD69, PD1 and OX40 were significantly increased in Tc17 cells compared to IL-17-CD8+ cells (Fig.4A,B). Not only the frequency of Tc17 cells expressing these markers, but also the intensity of their expression on the cell (Fig.4C) was increased. This does not simply reflect activation conditions, as the IL-17-cells were producing IFN-γ and were exposed to the same anti-CD3 / 28 stimulation as the Tc17 population.

**Figure 4:**
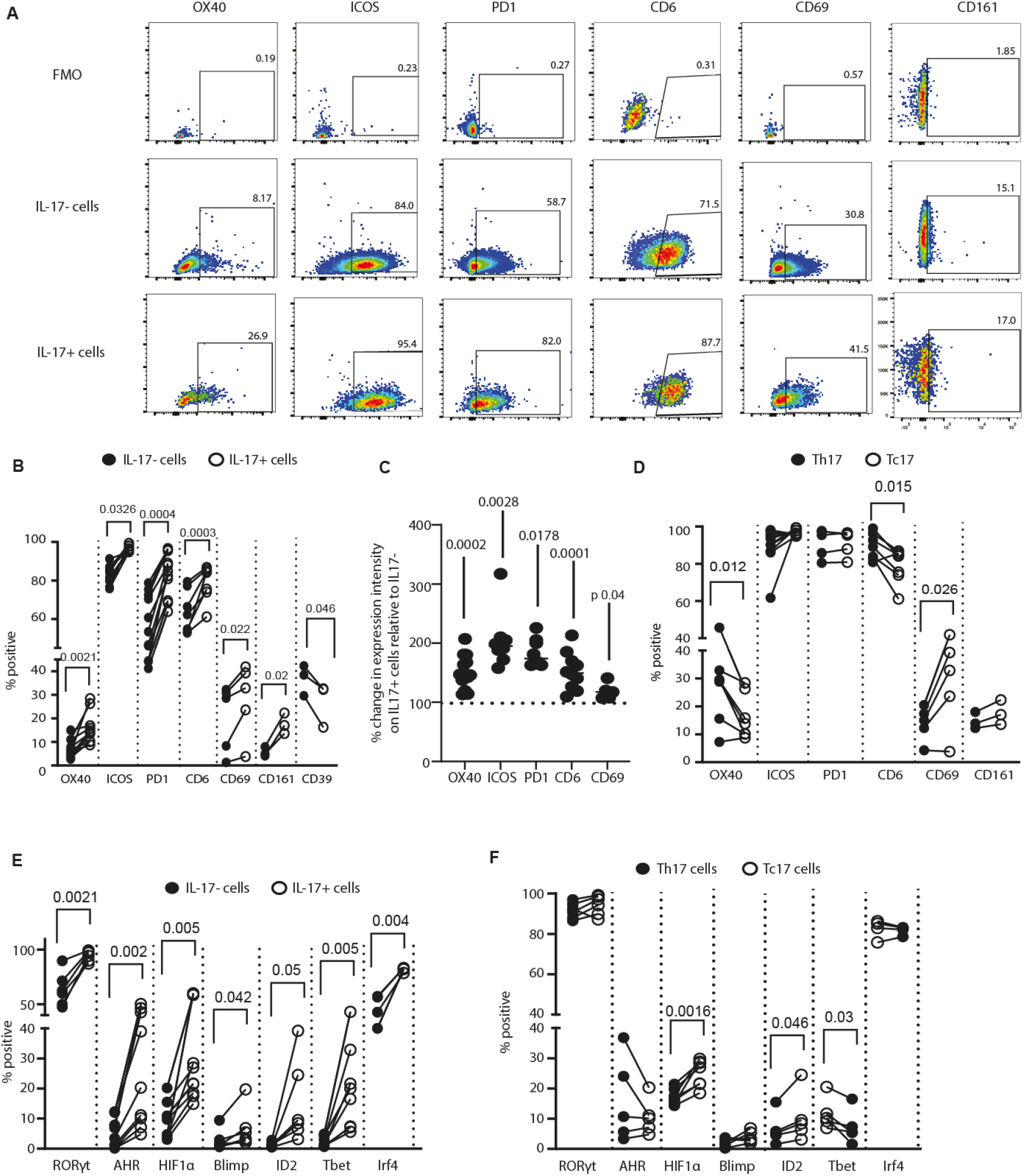
Tel7 cells express high surface levels of ICOS, PD1 and CD6. Splenocytes from C57BL/6J mice were cultured for 48 hours in Tel 7 driving conditions. Surface expression of (A-D) activation markers and (E-F) intracellular expression of transcription factors was assessed by flow cytometry. Data shown are individual mice with (C) line at median. Before-after analysis (B,D,E,F) compares IL-17- and IL-17+ T cells in the same experimental sample. Statistical significance was determined using (B, D, E, F) paired Student’s t tests; (C) paired Student’s t test on raw data before conversion. (B,D,E,F) Black symbols show IL-17-cellsand open symbols show IL-17+ cells. N values: B-D = 3-11; E-F = 6-9.

Importantly, the pattern of expression of ICOS, CD6, PD1, and CD69 were replicated in our culture conditions as they were in the *Tabula muris* dataset, validating that we are producing cells *in vitro* which substantially represent those found *in vivo*.

Although it was strongly increased compared to Tc1 cells, CD69 was only expressed on a minority of the Tc17 cells, and this was variable. The geometric mean was also only slightly increased. Similarly, OX40 was increased, but again only on a minority of cells, with an average of 15.8% of the Tc17 cells being OX40^+^. In contrast, ICOS, CD6 and PD1 were the strongest expressed (Fig4A,B,C). They were expressed on close to 100% of the Tc17 cells, and the geometric mean of ICOS and PD1 almost doubled compared to IL-17-cells (Fig.4C). Previous work has indicated CD39 and CD161 as being identifiers of human Tc17 cells. We found that murine cells did show substantially increased expression of CD161 compared to IL-17-cells, although at the 48-hour timepoint this was still on a minority of the cells; however, in contrast to human cells, CD39 was not increased (Fig.4B).

Next, we examined how the surface marker expression of Tc17 cells differs compared to their Th17 counterparts. OX40 and CD6 were expressed more strongly on Th17 cells than Tc17, but ICOS and PD1 were comparable in expression. Interestingly, CD69 was more strongly expressed on the Tc17 cell surface (Fig.4D). This is a different pattern of expression to that noted for human cells in which CD6 and CD69 were similarly expressed between Th17 and Tc17 cells but PD1 was altered.

### Transcription factor expression by Tc17 and Th17 cells

Transcription factor expression was also analysed in all these cell populations, informed by the TM dataset. As expected, almost all Tc17 cells expressed RORγt and IRF4 (Fig.4E) ^12,24^. It has previously been shown that Tc17 cells express the aryl hydrocarbon receptor, although unlike Th17 cells its centrality in their differentiation has not yet been demonstrated. Tc17 cells were significantly more likely to express AHR (Fig.4E) but expression was variable.

HIF-1α was also strongly up-regulated (Fig.4F); this is known to drive differentiation of IL-17-producing cells through its interaction with RORγt^35^. Interestingly, Id2 was very strongly expressed at the gene level, and was increased at the protein level in Tc17 cells but this remained only expressed by a minority of cells.

Tbet was not expressed on a high percentage of Tc17 cells, but unexpectedly was increased compared to IL-17-cells, which showed lower than expected expression. Unlike the other markers discussed here, in this case our culture results did not reflect the data from the *Tabula muris* analysis (Fig.2B, *Tbx21*) and so we believe this suppression of Tbet in Tc1 cells may be a consequence of the TGF-β in our culture ^36^.

### Metabolic alterations in Tc17 vs Th17 cells

In our analysis of the *Tabula muris* dataset we noted striking alterations in the expression of metabolism-related genes. In particular, Tc17 cells showed substantially higher expression of genes relating to metabolic stress adaptation, such as *Fabp5, Nnt, Oat* and *Adh1* than either IL-17^−^ CD8^+^ cells or Th17 cells (Fig.5A, text in red). In addition, there were a large number of differences between Tc17 and Th17 cells – in all cases, Tc17 cells had a lower level of this gene expression. These genes included *Acadl, Ass1, Ckmt1, Cox5a, Fasn, Gapdh, Pccb*, and *Slc2a1* (Fig.5A, text in blue). This gene expression pattern suggests a shift in Tc17 towards a glycolysis-supported metabolic state.

**Figure 5:**
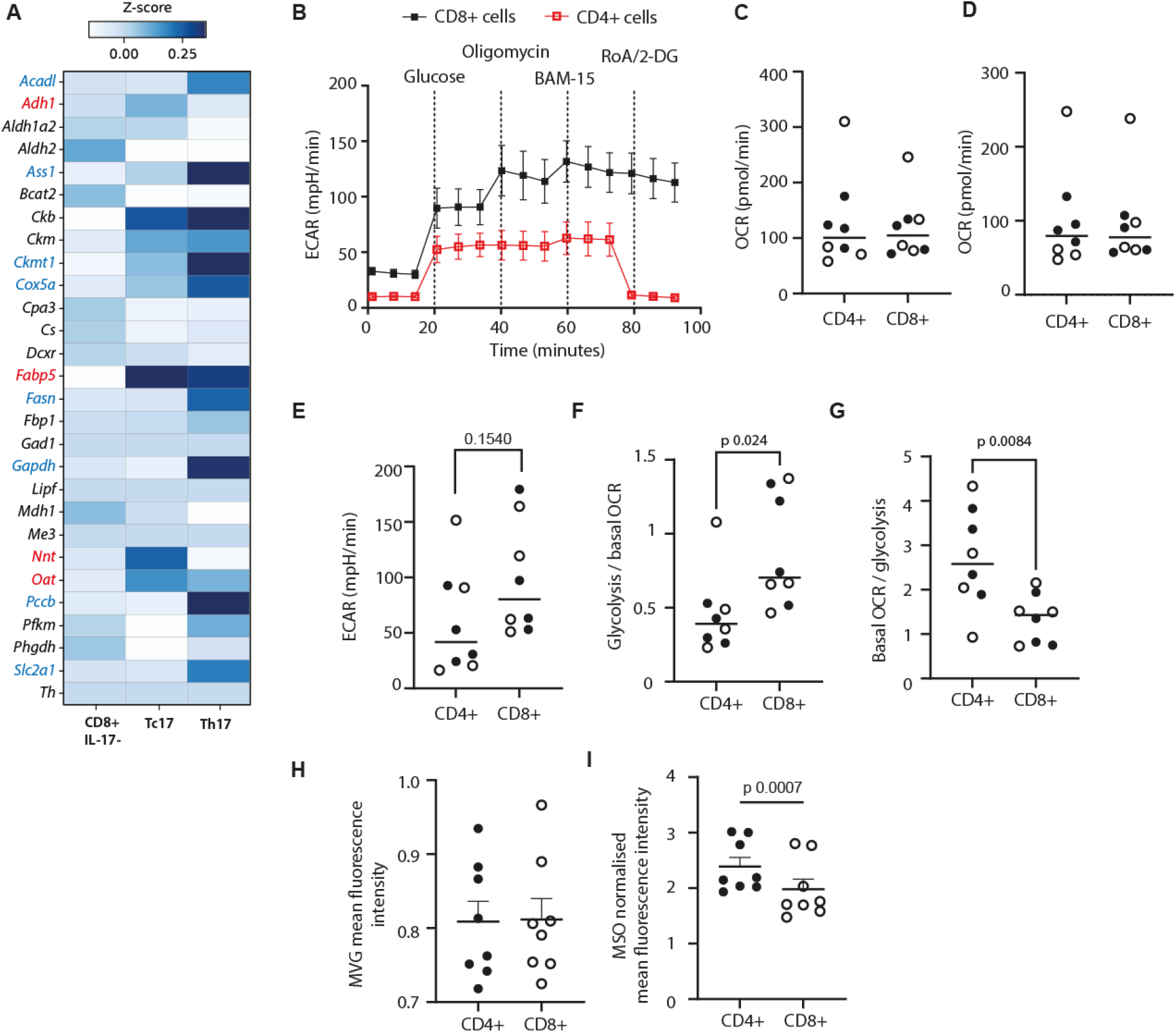
CD8+T cells in IL-17 driving conditions are more glycolytic than CD4+cells. (A) Analysis of the Tabula muris dataset identified metabolism-related genes differentially expressed between Tc1,Tc 17 and Th 17 cells. (B-G) Isolated T cells from CD57BL76J mice were analysed on a Seahorse metabolic extracellular flux analyser. (H) mitochondrial mass was assessed by incubation with mitoview green and (I) mitochondrial membrane potential was assessed by incubation with MitoSpy Orange, before both were analysed by flow cytometry. (C-G) filled circles represent male mice and open circles represent female mice. Statistical significance was assessed by paired Student’s t test. N values - 8 mice.

These alterations were intriguing. We therefore performed metabolic analyses on isolated CD4^+^ and CD8^+^ T cells in IL-17 driving conditions for 48 hours. Representative results from Seahorse assays are shown in Fig.5B.

The basal or maximal oxygen consumption rates were no different between CD8^+^ and CD4^+^ cells under these conditions (Fig.5C,D), in either male or female mice. The extracellular acidification rate was higher in CD8^+^ cells, although this was variable between the mice (Fig.5E). However, calculations from this data revealed that the glycolysis / basal OCR rate was significantly higher in CD8^+^ cells (Fig.5F), with therefore basal OCR / glycolysis rate being significantly lower (Fig.5G).

Flow cytometric analysis of mitochondrial phenotype in these cell types was then performed. The mitochondrial mass, measured by MitoView Green assay, was comparable between the cells (Fig.5H), but the membrane potential, measured by MitoSpy Orange, was significantly different (Fig.5I). CD8^+^ T cells had lower mitochondrial membrane potential than CD4^+^ cells in these IL-17-driving conditions.

## Discussion

In this study we set out to establish the phenotype, cytokine production, differentiation pathways and metabolic profile of IL-17-producing CD8^+^ T cells, comparing all to both IL-17^−^ CD8^+^ cells and Th17 cells.

Firstly, we confirmed that TGF-β is required and IL-1β is not for the consistent differentiation of Tc17 cells *in vitro*. A number of papers have used IL-1β in their differentiation cocktail-however, we demonstrate that it is the presence of TGF-β which induces differentiation of a clear population of cells which is both RORγt and IL-17-expressing but negative for IFN-γ and granzyme B. While other work using CD4^+^ T cells has shown IL-1β induces IL-17A alone, and TGF-β induces A/F co-expressing cells^37^, we do not find this to be the case in our studies. Here, IL-1β over a 96-hour culture led to a far lower production of IL-17A compared to TGF-β.

It is possible that this is an inherent difference between Th17 and Tc17 cells, with Tc17s being more disposed to producing IL-17F. Throughout our analyses in our *in vitro* culture of spleen cells, CD8^+^ T cells which produced either IL-17A or IL-17F tended to produce more of the latter than the former. However, spleen cells isolated and stained directly *ex vivo* produce more IL-I7A than IL-17F. This difference may suggest that the baseline for Tc17 cells is to produce more IL-17A, but that during inflammation or in the presence of high concentrations of cytokines, they switch on high production of IL-17F. Previous work has demonstrated that pathogenic Tc17 cells in the spinal cord during experimental autoimmune encephalomyelitis express IL-17A, with IL-17F suggested to be produced only by cells at mucosal sites^37^. Whether our *in vitro-*generated Tc17s therefore reflect mucosal Tc17 in surface marker expression or functional output remains to be examined.

These cells also produce cytokines other than the IL-17 family. Analysis of human Tc17 in the intestine demonstrated that 53% are co-producers of IFN-γ, which is in contrast to our cells generated in culture. This may be a consequence of *in vitro* versus *in vivo* stimulation, or the chronic extensive inflammation occurring in the intestine of Crohn’s patients triggering Tc17 cells to pick up Tc1 phenotypes, as Th17 cells do in long term inflammation ^38,39^. Certainly, the lack of IFN-γ that we see in cultured Tc17 cells reflects its absence in Th17 cells also.

Otherwise, the cytokine production by cultured Tc17 cells recapitulated that seen in the *Tabula muris* dataset, with increased production of IL-22 and IL-2 and a lack of IL-21. TNF was also universally produced. The increased production of IL-2 is very interesting, as Th17 cells show a suppression of the *Il2* gene, and indeed here we found Th17 cells did not produce IL-2 at all. How this gene is differentially regulated in Tc17 vs Th17 cells will be important to establish.

We also noted consistent alterations in cell surface marker expression in Tc17 cells compared to Tc1 cells, including almost universal expression of CD6, ICOS, and PD1 as well as CCR6 and strongly up-regulated CD161. Importantly, the pattern of expression that we noted in culture repeated what we found expressed in the public *Tabula muris* dataset, again confirming that our conditions are in general physiologically representative. PD1 inhibits the Tc17 to Tc1 transition, and to maintain the cells in their IL-17-producing phenotype ^40^. The increased expression of OX40 was particularly intriguing. The function of OX40 on these cells is unclear, as in CD4^+^ cells its triggering suppresses IL-17 production through chromatin remodelling ^41^. In humans, an OX40^hi^ PD1^lo^ Tc17 cell type has been detected in patients with human distal bile duct cancer ^21^, where its presence correlates with reduced survival. In the cells we generated in culture, high expression of OX40 was not accompanied by low expression of PD1 – this combination may be unique to pathological chronic conditions, and will need more mechanistic investigation to unravel, and to understand what triggers drive expression of OX40 on some Tc17 cells and not others, and whether these have different roles *in vivo*.

Metabolic differences were observed between CD4^+^ and CD8^+^ cells in their response to IL-17-driving conditions. The cells showed similar basal and maximal respiration, but CD8^+^ cells had reduced mitochondrial membrane potential and increased ECAR versus CD4^+^ cells, consistent with increased glycolytic lean. Transcriptome analysis of the *Tabula muris* dataset showed up-regulation of lipid-handing and redox-associated genes (*Fabp5, Nnt, Oat, Adh1)* as well as down-regulation of fatty acid oxidation and ETC-assocated genes (*Acadl, Pccb, Ass1, Cox5a)*.

This work agrees with previous findings^42^ that activated CD8^+^ T cells are less oxidative than their CD4^+^ counterparts in non-T cell-skewing culture conditions. We find this glycolytic lean to be maintained in IL-17-driving conditions.

Together, these data demonstrate firstly that culture conditions established here (48 hours with TGF-β, IL-6 and IL-23) generate large numbers of Tc17 cells that substantially mimic those found *in vivo*. We show that Tc17 cells can be identified through their expression of CD6, ICOS, PD1 and CD161. We also demonstrate phenotypic differences between Th17 and Tc17 cells, with the CD8+ cells being more glycolytic and producing far more IL-2, IL-22 and IL-17F. These findings lend weight to the growing opinion that Th17 and Tc17 cells have differential roles in homeostasis and inflammation.

## Methods

### Mice

Wild-type C57Bl/6JOlaHsd mice were bred and housed in individually ventilated cages, under specific pathogen-free conditions. Male and female mice between 6-12 weeks of age were used. Mice were kept in environmental conditions in line with the ASPA Code of Practice for the United Kingdom – temperatures between 19 and 24^°^C, humidity between 45 and 65%, and 12 hours of light then 12 hours of dark. All animal experiments were performed by fully trained personnel in accordance with Home Office UK project licence PAF438439.

### Murine tissue and single-cell preparations

Single-cell preparations of splenocytes were achieved by mechanical disruption through a 100 μM strainer and washing with PBS. Red blood cells were lysed using RBC Lysis Buffer, as per the manufacturer’s instructions (BD Biosciences, #555899).

### In vitro T cell subset differentiation

200,000 splenocytes were plated per well in round-bottom 96-well plates in complete medium (RPMI, 10% foetal calf serum, 10 units/ml penicillin, 10 μg/ml streptomycin and 2 mM L-glutamine, all supplied by Gibco, ThermoFisher UK). Cells were differentiated in the presence of plate-bound anti-CD3 (5 μg/ml; Biolegend, #100339) in the presence or absence of αCD28 (1 μg/ml; Biolegend, #102115), with a combination of recombinant cytokines - IL-6 (50 ng/ml; Biolegend, #575706), IL-23 (50 ng/ml; Biolegend, #589006) TGFβ1 (10 ng/ml; Biolegend, #580706), IL-23 (50 ng/ml; Biolegend, #589006) or IL-1β (10 ng/ml, Biolegend, # 575102).

### Flow cytometry

Cells were incubated for 4 hours at 37°C with Cell Stimulation Cocktail containing protein transport inhibitors (ThermoFisher, #00-4970-03) prior to staining for cellular viabilty using LIVE/DEAD™ Fixable Yellow Dead Cell Stain Kit (405 nm excitation, ThermoFisher Scientific, Invitrogen # L34959) in PBS for 20 minutes at room temperature. Cells were then stained for surface markers (refer to the antibodies section for dilutions) in 2% foetal bovine serum PBS, for 30 minutes at 4 °C, protected from light. Samples were then fixed and permeabilised using BD Cytofix/Cytoperm (BD Biosciences, #554722), as per the manufacturer’s guidelines (refer to antibodies section for dilutions), before being stained with antibodies detecting cytokines or transcription factors for 30 minutes at 4 °C, protected from light.

**Table 1.** Antibody reagents used for flow cytometry.

| Marker | Fluorophore | Clone | Supplier | Catalogue | Dilution |
| --- | --- | --- | --- | --- | --- |
| *Surface markers* |  |  |  |  |  |
| CD4 | PerCPCy5.5 | GK1.5 | BioLegend | 100434 | 1:200 |
| CD5 | Pacific Blue | 53-7.3 | BioLegend | 100642 | 1:150 |
| CD6 | PE | OX-129 | BioLegend | 146403 | 1:150 |
| CD8a | PE | 53-6.7 | BD | 100708 | 1:200 |
| CD8a | APC-Fire 750 | 53-6.7 | BD | 100766 | 1:200 |
| CD69 | AF700 | H1.2F3 | BioLegend | 104539 | 1:150 |
| CD134 (OX40) | APC | OX-86 | BioLegend | 119414 | 1:150 |
| CD278 (ICOS) | PE-Cyanine7 | C398.4A | BioLegend | 313520 | 1:200 |
| CD279 (PD-1) | APC | 29F.1A12 | BioLegend | 135240 | 1:200 |
| *Cytokine markers* |  |  |  |  |  |
| IFNγ | FITC | XMG1.2 | BioLegend | 505806 | 1:100 |
| IL-17A | PE/Dazzle 594 | TC11-18H10.1 | BioLegend | 506938 | 1:150 |
| IL-17F | AF647 | 9D3.1C8 | BioLegend | 517004 | 1:150 |
| IL-17F | AF488 | 9D3.1C8 | BioLegend | 517006 | 1:150 |
| *Transcription factors* |  |  |  |  |  |
| AHR | PE-Cyanine7 | 4MEJJ | eBiosciences | 25-5925-80 | 1:100 |
| Blimp | APC | 29F.1A12 | BioLegend | 135210 | 1:200 |
| HIF1α | PE | Mgc3 | ThermoFisher | 12-7528-82 | 1:150 |
| ID2 | eFluor450 | ILCID2 | ThermoFisher | 48-9475-82 | 1:100 |
| RORγT | PE | B2D | ThermoFisher | 12-6981-82 | 1:100 |
| TBet | PE-Cyanine7 | 4B10 | BioLegend | 644824 | 1:100 |

Samples were collected using an BD FACSAriaII cytometer (BD Biosciences) and Diva software version 9, and analysed using FlowJo software version 10 (BD Biosciences).

### Analysis of Tabula muris dataset

We analysed the complete *Tabula Muris* dataset, collected in March 2025, which contains mouse transcriptomic data for 356,213 cells across 20 organs ^32^. Cells expressing either CD8α or CD8β1, and either Il17a or Il17f were classified as Tc17 cells, whereas cell expressing CD4 and either Il17a or Il17f were classified as Th17 cells. Raw gene counts were normalised, log-transformed, and scaled using Scanpy ^43^ in Python.

### Extracellular flux analysis

Analysis of the OCR (pmol/min) and ECAR (mpH/min) was conducted using the Seahorse XFe96 metabolic extracellular flux analyser (Agilent). CD4^+^ or CD8^+^ T cells were isolated from whole splenocytes using positive magnetic selection and incubated with IL-17-polarising cytokines as previously, for 48 hours. T cells were then harvested and resuspended in serum-free unbuffered RPMI 1640 medium, before being plated onto Seahorse cell plates (2.5×10^5^ cells per well) coated with poly-d-lysine (Invitrogen) to enhance T cell adhesion. Profiling was conducted using the addition of glucose (10mM), oligomycin (1µM), Bam-15 (3µM) and 2-DG/rotenone/antimycin A (2-DG at 50mM, rotenone and antimycin both at 2µM, all are given as final concentrations, all are from Sigma-Aldrich) which are injected in during the course of the assay.

### Flow cytometric analysis - metabolism

To assess the mitochondrial mass of the cells, following cell culturing in the conditions mentioned previously, cells were incubated in RPMI with MitoView Green (50 nM; Biotium, Cat#70054) for 20 mins at 37°C and 5% CO_2_ prior to washing and analysis. To assess the mitochondrial membrane potential (ΔΨm) cells were incubated in RPMI with MitoSpy Orange (MSO, 25nM, Biolegend Cat#424804) for 20 min at 37°C and 5% CO_2_ ± the mitochondrial uncoupler, Bam-15 (3 μM, BioTechne, Cat#5737) prior to washing and analysis. Addition of Bam-15 dissipates mitochondrial membrane potential permitting measurement of baseline MSO staining in depolarised mitochondria. Data are presented as a ratio of MSO MFI in absence of Bam-15/ MSO MFI in presence of Bam-15. Following incubation with the Mitotracker probes, samples were transferred to FACS tubes and acquired using flow cytometry as previously.

### Statistics

All data shown are expressed as individual data points with line at median. Analysis was performed with GraphPad Prism software. A minimum of three mice was used for *in vitro* experiments, in individually performed experiments as indicated in figure legends. Multiple groups were compared by two-way analysis of variance tests with either Tukey’s multiple comparisons tests. Statistical differences between the groups were determined by two-way ANOVA. p<0.001 is symbolized by ***; p<0.01 by **; and p<0.05 by *.

## Data availability

The datasets generated during the current study are available from the corresponding author (EGF) upon request. The authors declare that all data supporting the findings of this study are available within the paper.

## References

1. Langrish, C. L. et al. IL-23 drives a pathogenic T cell population that induces autoimmune inflammation. J. Exp. Med. 201, 233–240 (2005).

2. Harrington, L. E. et al. Interleukin 17–producing CD4+ effector T cells develop via a lineage distinct from the T helper type 1 and 2 lineages. Nat. Immunol. 6, 1123–1132 (2005).

3. Veldhoen, M., Hocking, R. J., Atkins, C. J., Locksley, R. M. & Stockinger, B. TGFβ in the Context of an Inflammatory Cytokine Milieu Supports De Novo Differentiation of IL-17-Producing T Cells. Immunity 24, 179–189 (2006).

4. Schnell, A., Littman, D. R. & Kuchroo, V. K. TH17 cell heterogeneity and its role in tissue inflammation. Nat. Immunol. 24, 19–29 (2023).

5. Papadopoulou, G. & Xanthou, G. Metabolic rewiring: a new master of Th17 cell plasticity and heterogeneity. FEBS J. 289, 2448–2466 (2022).

6. Park, E. & Ciofani, M. Th17 cell pathogenicity in autoimmune disease. Exp. Mol. Med. 57, 1913–1927 (2025).

7. Hamada, H. et al. Tc17, a unique subset of CD8 T cells that can protect against lethal influenza challenge. J. Immunol. Baltim. Md 1950 182, 3469–3481 (2009).

8. Ge, C. et al. Mouse CD8+ T cell subsets differentially generate IL-17-expressing cells in the colon epithelium and lamina propria. Clin. Exp. Immunol. 219, uxae120 (2025).

9. Globig, A.-M. et al. High-dimensional profiling reveals Tc17 cell enrichment in active Crohn’s disease and identifies a potentially targetable signature. Nat. Commun. 13, 3688 (2022).

10. Ciric, B., El-behi, M., Cabrera, R., Zhang, G.-X. & Rostami, A. IL-23 drives pathogenic IL-17-producing CD8+ T cells. J. Immunol. Baltim. Md 1950 182, 5296–5305 (2009).

11. Mills, K. H. G. IL-17 and IL-17-producing cells in protection versus pathology. Nat. Rev. Immunol. 23, 38–54 (2023).

12. Huber, M. et al. IL-17A secretion by CD8^+^ T cells supports Th17-mediated autoimmune encephalomyelitis. J. Clin. Invest. 123, 247–260 (2013).

13. Peelen, E. et al. Fraction of IL-10+ and IL-17+ CD8 T cells is increased in MS patients in remission and during a relapse, but is not influenced by immune modulators. J. Neuroimmunol. 258, 77–84 (2013).

14. Interleukin-17+CD8+ T Cells Are Enriched in the Joints of Patients With Psoriatic Arthritis and Correlate With Disease Activity and Joint Damage Progression - Menon - 2014 - Arthritis & Rheumatology - Wiley Online Library. https://acrjournals.onlinelibrary.wiley.com/doi/10.1002/art.38376 (2024).

15. Wang, H. H. et al. Interleukin-17-secreting T cells in neuromyelitis optica and multiple sclerosis during relapse. J. Clin. Neurosci. 18, 1313–1317 (2011).

16. Tom, M. R. et al. Novel CD8+ T-Cell Subsets Demonstrating Plasticity in Patients with Inflammatory Bowel Disease. Inflamm. Bowel Dis. 22, 1596–1608 (2016).

17. Smillie, C. S. et al. Intra- and Inter-cellular Rewiring of the Human Colon during Ulcerative Colitis. Cell 178, 714-730.e22 (2019).

18. Saito, H., Yamada, Y., Takaya, S., Osaki, T. & Ikeguchi, M. Clinical relevance of the number of interleukin-17-producing CD 8+ T cells in patients with gastric cancer. Surg. Today 45, 1429–1435 (2015).

19. Lee, Y. H. et al. IFNγ−IL-17+ CD8 T cells contribute to immunosuppression and tumor progression in human hepatocellular carcinoma. Cancer Lett. 552, 215977 (2023).

20. Kuang, D.-M. et al. Tumor-Activated Monocytes Promote Expansion of IL-17–Producing CD8+ T Cells in Hepatocellular Carcinoma Patients. J. Immunol. 185, 1544–1549 (2010).

21. Chellappa, S. et al. CD8+ T Cells That Coexpress RORγt and T-bet Are Functionally Impaired and Expand in Patients with Distal Bile Duct Cancer. J. Immunol. Baltim. Md 1950 198, 1729–1739 (2017).

22. He, D. et al. CD8+ IL-17 producing T cells are important in effector functions for the elicitation of contact hypersensitivity responses. J. Immunol. Baltim. Md 1950 177, 6852–6858 (2006).

23. Liu, H.-P. et al. TGF-β converts Th1 cells into Th17 cells through stimulation of Runx1 expression. Eur. J. Immunol. 45, 1010–1018 (2015).

24. Yen, H.-R. et al. Tc17 CD8 T Cells: Functional Plasticity and Subset Diversity. J. Immunol. Baltim. Md 1950 183, 7161–7168 (2009).

25. Arra, A. et al. The differentiation and plasticity of Tc17 cells are regulated by CTLA-4-mediated effects on STATs. OncoImmunology 6, e1273300 (2017).

26. Erice, P. A. et al. Downregulation of Mirlet7 miRNA family promotes Tc17 differentiation and emphysema via de-repression of RORγt. eLife 13, RP92879 (2024).

27. Lin, G. et al. The Pro-inflammatory Functions of Type 1 CD 8+ T Cells and Interleukin-17-producing Cluster of Differentiation 8+ T Cells are Exhausted by Cholesterol in Atherosclerosis. Iran. J. Immunol. 21, 353–364 (2024).

28. Minns, D. et al. The neutrophil antimicrobial peptide cathelicidin promotes Th17 differentiation. Nat. Commun. 12, 1285 (2021).

29. Huber, M. et al. A Th17-like developmental process leads to CD8+ Tc17 cells with reduced cytotoxic activity. Eur. J. Immunol. 39, 1716–1725 (2009).

30. Saxena, A. et al. Tc17 CD8+ T Cells Potentiate Th1-Mediated Autoimmune Diabetes in a Mouse Model. J. Immunol. 189, 3140–3149 (2012).

31. El-Behi, M. et al. Committed Tc17 cells are phenotypically and functionally resistant to the effects of IL-27. Eur. J. Immunol. 44, 3003–3014 (2014).

32. The Tabula Muris Consortium et al. Single-cell transcriptomics of 20 mouse organs creates a Tabula Muris. Nature 562, 367–372 (2018).

33. Giordano, M. et al. Molecular profiling of CD 8 T cells in autochthonous melanoma identifies Maf as driver of exhaustion. EMBO J. 34, 2042–2058 (2015).

34. Ponomarev, E. D. et al. GM-CSF Production by Autoreactive T Cells Is Required for the Activation of Microglial Cells and the Onset of Experimental Autoimmune Encephalomyelitis. J. Immunol. 178, 39–48 (2007).

35. Dang, E. V. et al. Control of TH17/Treg Balance by Hypoxia-Inducible Factor 1. Cell 146, 772–784 (2011).

36. Park, I.-K., Shultz, L. D., Letterio, J. J. & Gorham, J. D. TGF-β1 Inhibits T-bet Induction by IFN-γ in Murine CD4+ T Cells through the Protein Tyrosine Phosphatase Src Homology Region 2 Domain-Containing Phosphatase-1. J. Immunol. 175, 5666–5674 (2005).

37. Wanke, F. et al. Expression of IL-17F is associated with non-pathogenic Th17 cells. J. Mol. Med. 96, 819–829 (2018).

38. Smith, K. J. et al. The antimicrobial peptide cathelicidin drives development of experimental autoimmune encephalomyelitis in mice by affecting Th17 differentiation. PLoS Biol. 20, e3001554 (2022).

39. Hirota, K. et al. Fate mapping of IL-17-producing T cells in inflammatory responses. Nat. Immunol. 12, 255–263 (2011).

40. Arra, A., Lingel, H., Pierau, M. & Brunner-Weinzierl, M. C. PD-1 limits differentiation and plasticity of Tc17 cells. Front. Immunol. 14, (2023).

41. Xiao, X. et al. The Costimulatory Receptor OX40 Inhibits Interleukin-17 Expression through Activation of Repressive Chromatin Remodeling Pathways. Immunity 44, 1271–1283 (2016).

42. Cao, Y., Rathmell, J. C. & Macintyre, A. N. Metabolic Reprogramming towards Aerobic Glycolysis Correlates with Greater Proliferative Ability and Resistance to Metabolic Inhibition in CD8 versus CD4 T Cells. PLoS ONE 9, e104104 (2014).

43. Wolf, F. A., Angerer, P. & Theis, F. J. SCANPY: large-scale single-cell gene expression data analysis. Genome Biol. 19, 15 (2018).

